# Exploring the uncharted: Novel potential filariasis vectors unveiled in Sri Lanka

**DOI:** 10.1101/2024.08.23.609308

**Authors:** Sachini U. Nimalrathna, V.A.Chathuri B. Amarasiri, D. Kaveesha R. Aluthge, Chandana H. Mallawarachchi, T.G.A. Nilmini Chandrasena, B. G. D. Nissanka K. de Silva, Nilanthi R. de Silva, Michael J. Kimber, Hiruni Harischandra

## Abstract

Brugian filariasis (BF) has reemerged in Sri Lanka recently. Studies suggest the emergence of a variant brugian parasite. Knowledge on transmission dynamics is important in restraining the spread of infection. This study investigated the potential vector mosquitoes of this variant brugian parasite around six indexed human BF cases in five BF endemic districts in Sri Lanka. A total of 1711 mosquitoes from 20 species were analyzed. Potential infective mosquitoes were detected by the presence of L_3_ larval stage of brugian parasites within the head and thorax regions upon dissections and confirmed by amplification of the *Brugia* species-specific *HhaI* region. Twelve (12) mosquito species that could potentially serve as vectors for BF transmission in selected endemic areas in the country were identified due to the presence of L_3_ larvae in the head and thorax regions. This is the first report of *Ma. indiana, Ar. subalbatus*, *Ae. albopictus, Cq. crassipes, Cx. tritaeniorhynchus, Cx. bitaeniorhynchus, Cx. quinquefasciatus*, *Cx. gelidus, Cx. lopoceraomyia and Cx. vishnui* with the potential of serving as vectors for BF transmission in Sri Lanka and *Cx. bitaeniorhynchus*, *Cx. gelidus, Cx. lopoceraomyia and Cx. vishnui* in the world through a field study. Of these, *Ma. indiana, Cx. tritaeniorhynchus, Cx. quinquefasciatus* and *Ar. subalbatus* together with *Ma. uniformis, Ma. annulifera* had the highest prevalence and infection rate at certain study sites. The recovery of parasite-positive *Ma. indiana, Cx. quinquefasciatus* and *Ar. subalbatus* mosquitoes from window traps demonstrate the probable anthropophilic nature of these species, and their high prevalence in the respective areas deems further investigation on their vector potentiality essential.

**Author Summary:** Lymphatic filariasis (LF) is a debilitating disease affecting over 120 million people worldwide. Of the two types of LF, Brugian Filariasis (BF) has a wide range of definitive hosts, making it possible for the rapid spread of the disease. BF was considered eliminated from Sri Lanka in 1969 but is on the rise after four decades of quiescence. Effective vector control is an integral part of infection control. We carried out entomological investigations for potential vectors of re-emerged BF in five BF-endemic areas in Sri Lanka. Here we report *Ma. indiana, Cx. tritaeniorhynchus, Cx. bitaeniorhynchus, Cx. quinquefasciatus*, *Cx. gelidus, Cx. lopoceraomyia, Cx. vishnui, Ar. subalbatus*, *Ae. albopictus* and *Cq. crassipes* with the potential to serve as vectors for BF transmission in Sri Lanka due to the presence of L_3_ larvae in the head and thorax regions and *Cx. bitaeniorhynchus, Cx. quinquefasciatus*, *Cx. gelidus, Cx. lopoceraomyia and Cx. vishnui* in the world. The high prevalence and infection rate of *Ma. indiana, Cx. quinquefasciatus* and *Ar. subalbatus* mosquitoes and their anthropophagic nature in certain BF endemic areas are worrisome. This study emphasizes the importance of a comprehensive analysis of BF vectors for vector control strategies tailored to each region and season.

## Introduction

Lymphatic filariasis (LF) is a mosquito-borne, Neglected Tropical Disease (NTD) caused mainly by the four filarial species, *Wuchereria bancrofti*, *Brugia malayi*, *Brugia pahangi* and *Brugia. timori*. Worldwide, 859 million people are at risk in 50 endemic countries, including Sri Lanka (1). The most recognizable symptoms of LF are hydrocele and lymphoedema; which can lead to elephantiasis that causes massive swelling of the extremities. These symptoms are attributed to the inflammatory reaction caused by the adult parasites residing within the lymphatic vasculature leading to lymphatic vessel damage and dysfunction which predispose lymphedema of the lower limbs. This makes LF one of the leading causes of permanent disfigurement and the second leading cause of long-term disability (2,3). Prior to the LF elimination programme, one-tenth of the inhabitants of the endemic region in Sri Lanka were at risk of being affected (4–7). The severe morbidity of this disease is a major socioeconomic burden and hinders the development of affected communities who reside in developing countries (8–12). The estimated economic burden is over 5.8 billion USD per year, which includes treatment, cost of healthcare and potential income loss (13).

In the past, both bancroftian and BF (nocturnal periodic strain) prevailed in the country for many years, bancroftian filariasis along the southwestern coastal belt known as the “filariasis belt” with scattered foci of BF in the endemic belt and towards the interior of the country (14). The National Programme for Elimination of LF (NPELF) in 2002 recognized three provinces, Western, Northwestern and Southern as endemic for bancroftian filariasis while BF was considered eliminated as cases were not reported since the late sixties. The elimination was attributed to the clearance of the aquatic vegetation required for the breeding of vector mosquitoes, *Mansonia* spp. (15). However, surveillance activities carried out following the mass chemotherapy programme reported sporadic occurrences of BF in all three LF-endemic provinces (16,17). The reemerged BF shows evidence of spreading and establishing itself in the endemic region, as surveillance in 2023 records an increase of BF cases over those of bancroftian filariasis (Personal communication). This situation calls for urgent action as it poses a serious challenge to the maintenance of the LF elimination as a public health problem status in the country. Unlike bancroftian filariasis, BF has a substantial zoonotic reservoir of infection, and therefore, it is imperative that a multipronged approach is implemented targeting, the parasites and the vectors.

BF is a vector-borne disease; therefore, vector control is an effective method of hindering transmission of the disease. A wide vector range for zoonotic and anthroponotic brugian parasites has been reported from many regions of the world, including *Anopheles, Culex, Aedes* and *Mansonia* (18). Distribution and transmission potential of these vectors vary greatly based on geography. In Sri Lanka, *Mansonia annulifera and Mansonia uniformis* have been implicated as zooanthropophagic vector mosquitoes for the transmission of periodic *B. malayi* in the past (19–21). However, a comprehensive analysis of vectors of the brugian parasite in Sri Lanka has not been conducted to date and is vital to implement effective, tailored control strategies for the elimination of the disease from the country.

This study investigated the natural infection of mosquitoes with the re-emerged *B. malayi* parasite. We report that *Mansonia indiana,* many *Culex* species*, Armigeres subalbatus*, *Aedes albopictus* and *Coquillettidia crassipes* showed evidence of infection and parasite development suggesting a potential vectorial capacity for the brugian parasites in Sri Lanka. Interestingly, *Culex bitaeniorhynchus, Culex quinquefasciatus*, *Culex gelidus, Culex lopoceraomyia* and *Culex vishnui* also showed evidence of natural infection and development of brugian parasites in Sri Lanka and have not been reported so previously from other BF endemic regions in the world. These findings suggest the importance of further entomological studies to detect mosquitoes supporting the development of filarial parasite to L_3_ stage (infective stage), the percentage of larvae reaching maturity and the anthropophilic nature of the mosquito for it to be an efficient vector.

## Methodology

### Study area selection and mosquito collection

Five districts along the filariasis belt of Sri Lanka with the highest number of human BF cases (Puttalam-52, Kalutara-32, Gampaha-23, Galle-10, Colombo-6) reported to the Anti-filariasis Campaign (AFC) by April 2021 were selected for the study. The most recently reported case in each district at the time of the study is referred to as the index case hereafter. A 500 m radius was mapped around the index case and was demarcated as the study site. This buffer zone was chosen to encompass the mean flight range of *Mansonia* spp. mosquitoes (350 m), the known vector for *B. malayi*. Two study sites were selected from the Puttalam district and one each from the other four districts. The selected study sites were Puruduwella-Puttalam (S_1_), Maggona-Kalutara (S_2_), Wattala-Gampaha (S_3_), Induruwa-Galle (S_4_), Boralesgamuwa-Colombo (S_5_) and Mahawewa-Puttalam (S_6_) (**Fig 1).**

**Fig 1:**
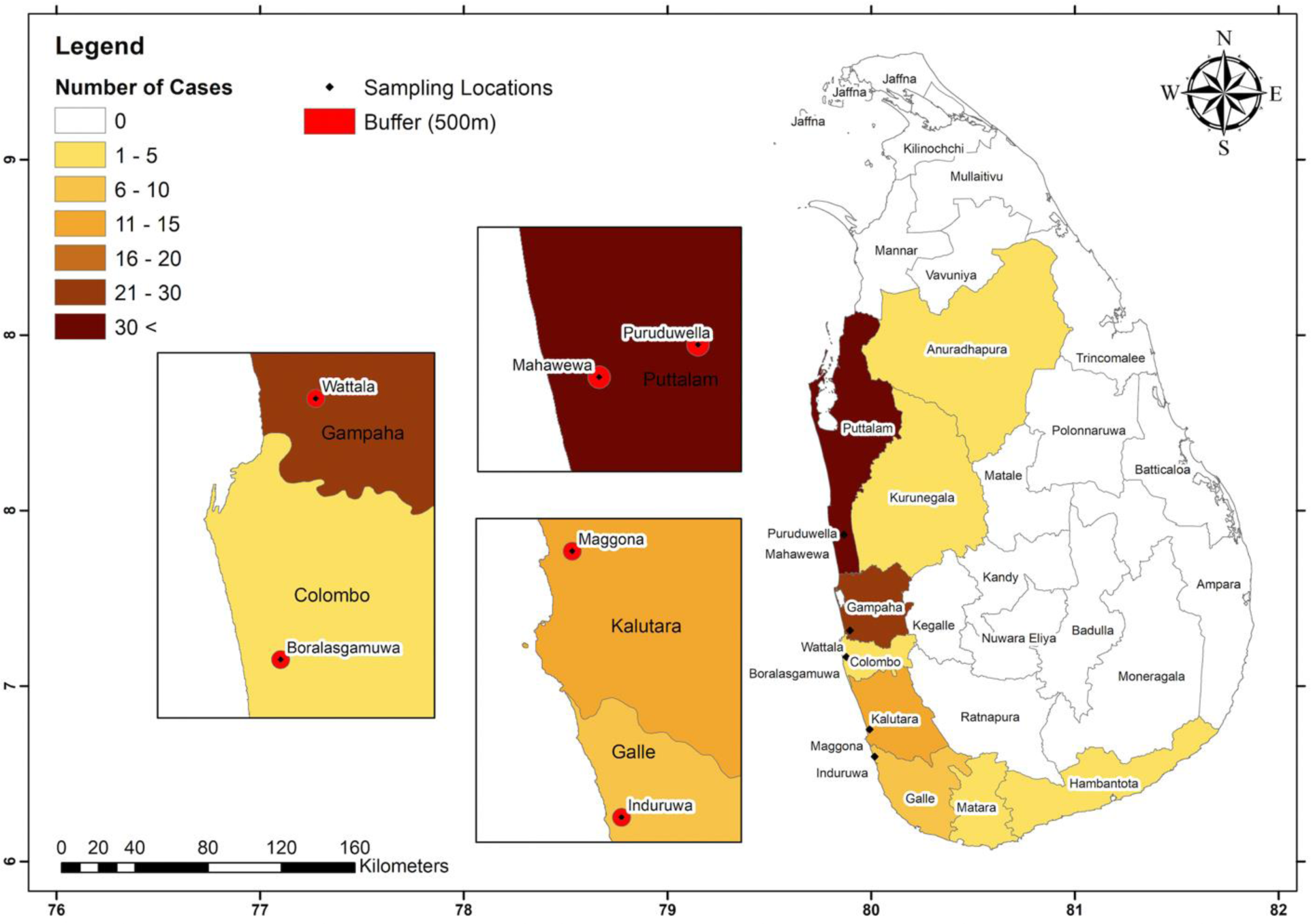
Study site selection. LF has been reported from nine districts in Sri Lanka. LF is endemic in all these districts except Anuradhapura, and this region is known as the filariasis belt. Five districts with the highest number of BF cases by April 2021 were selected (Puttalam-53, Kalutara-32, Gampaha-23, Galle-10, Colombo-6) for this study. Locations of the most recent BF cases reported from the selected districts were mapped and a 500 m buffer zone (depicted in red) around each case was designated as a study site (Puruduwella, Puttalam (S_1_), Maggona, Kalutara (S_2_) and Wattala, Gampaha (S_3_), Induruwa-Galle (S_4_), Boralesgamuwa-Colombo (S_5_) and Mahawewa-Puttalam (S_6_)).

### Detection of mosquito species that can serve as potential vectors for the brugian parasite prevalent in Sri Lanka

#### a. Mosquito sampling

Mosquitoes were collected during the monsoon season at each study site to identify the potential vectors of BF in Sri Lanka. Further sampling was done during the inter-monsoon season at the study areas in the three districts with the highest human disease incidence (S_1_, S_2_ and S_3_) to assess the effects of seasonal variation on the distribution of these vectors. Mosquitoes were collected between 18:00-04:00 h Sri Lankan Standard Time (SLST) using various traps to ensure that the mosquito collection was a true representation of the mosquito population at that time at each site. Dog-bait traps were assembled at the study site to attract zoophilic mosquitoes. A dog was tied overnight within a white, rectangular nylon mesh, tent-like trap (4 m × 3 m × 3 m) which was raised about 6 inches from the ground to allow mosquitoes to enter the trap. The following morning the dog was released, and the mosquitoes were collected using a mouth aspirator. Window traps of 56 cm X 56 cm X 46 cm made using standard procedures (22) were assembled on the outside of the windows to collect the anthropophilic mosquitoes exiting after a bloodmeal. The rest of the window was obstructed to ensure mosquitoes exited only through the opening leading to the trap. The trap was left in place all night, and mouth aspirators were used to collect the mosquitoes the following morning. Gravid traps were used to attract gravid mosquitoes in the area. An infusion of water and organic material (300g fresh cow manure, 150g of Gliricidia leaves, 10 g of yeast and 5 L of water) fermented for 1-4 weeks was placed in a plastic dishpan of 20 cm X 39 cm X 32 cm with a motorized suction trap and left overnight. The mosquito-infested net was carefully removed from the trap the following morning and mosquitoes were collected using a mouth aspirator. The species of all the collected mosquitoes were identified using standard morphological keys (23).

### Analysis of the spatial distribution of potential vectors within a study site

Additional mosquito collections were done at S_1_ to analyze the spatial distribution of infective vectors in relation to the index case. The most recent human case of BF was reported from S_1_, which is also the district with the highest number of human BF incidences. Mosquitoes were collected from the index site, approximately 150 m, 450 m and 650 m away from the index site **(Fig 2)** using dog-bait traps, window traps and gravid traps between 18:00-04:00 h SLST as before.

**Fig 2:**
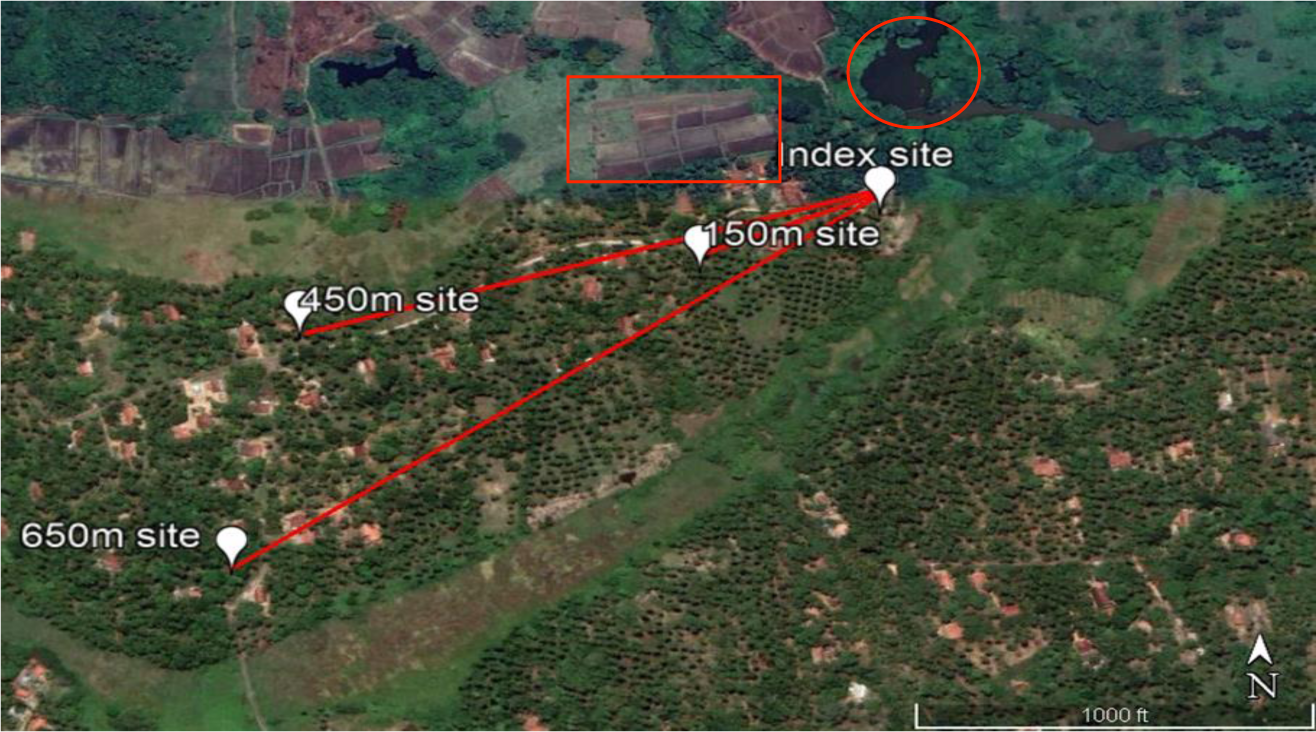
Study sites for the analysis of the spatial distribution of potential vectors. Mosquitoes were collected from the index site and 150 m, 450 m, and 650 m away from the index site at S_1_. Circled and rectangular areas show the natural lake and paddy field respectively situated near the sampling sites.

#### b. Detection of infected and potentially infective mosquitoes

Head and thorax regions of female mosquitoes were dissected to identify mosquitoes harboring L_3_ infective and other larval stages of the parasites. Each mosquito was placed on a slide under a binocular dissecting microscope and the legs and wings were removed. The head and thorax regions were separated from the abdomen and placed in a separate drop of saline and dissected to identify potentially infective (with L_3_ larval stage of the parasite) and infected (L_1_, L_2_, and L_3_ larval stage of the parasite) mosquitoes. Heads, thoraces and abdomen regions of the nematode parasite-positive mosquitoes were stored separately at −20°C.

DNA was extracted from the head and thoraces, and abdomens separately of each mosquito using the Blood and Tissue DNA extraction kit (Qiagen, Germany) as per manufacturer’s guidelines with some modifications as follows; the samples were homogenized using a vortex for about 2 mins at 2 000 rpm (Scientific Industries/ SI-A546). Next 180 µl of buffer ATL and 20 µl of proteinase K was added to the sample and vortexed for 2 mins at 2 000 rpm speed. Next, 200 µl of Buffer AL was added and the samples were vortexed for 1 min at 2 000 rpm and incubated at 56°C for 10 mins. Then 200 µl of ethanol (90-100%) was added and the sample was vortexed for 1 min at 2 000 rpm. Sample mixtures were pipetted into DNeasy Mini spin columns placed in 2 ml collection tube and centrifuged (Universal centrifuge, Gemmy, PLC-036H) at ≥6,000 g (8,000 rpm) of the centrifuge for 1 min. After discarding the flow through, 500 µl of Buffer AW_1_ was added. The sample was centrifuged again at ≥6,000 g for 1 min and the flow through was discarded. Next, 500 µl of Buffer AW_2_ was added and the samples were incubated at room temperature for 3 mins and centrifuged again at ≥20,000 g (14,000 rpm) for 1 min. Finally, the DNA was eluted twice to the same tube by adding 20 μl and 15 μl of nuclease free water respectively in two subsequent steps and centrifuging at ≥6,000 g for 1 min.

The *Brugia* species-specific *Hha1* region was amplified from the DNA extracts of the head and thorax regions of dissection-positive mosquitoes as previously described (Partono 1994). Briefly primers 5’-GCGCATAAATTCATCAGC-3 ’(Forward) and 5’-GCGCAAAACTTAATTACAAAAGC-3 ’(Reverse) were used to amplify the 322 bp *Hha1* repeat region of *Brugia* species. The PCR amplification was performed in a 25 μl reaction volume containing 5.0 μl of 5x PCR buffer, 0.5 μl (0.2 mM) dNTP, and 10 μM of each primer, 1U *Taq* polymerase, and 2 μl of the extracted template DNA. The PCR procedure consisted of an initial denaturation step at 94°C for 5 mins followed by 35 cycles each of denaturation at 94°C for 1 min, annealing at 59°C for 1 min, extension at 72°C for 1 min, and final extension at 72°C for 10 mins. The products were visualized by electrophoresis on a 1.5% agarose gel.

Selected positive PCR products were sequenced at the DNA sequencing facility at Iowa State University, USA and Macrogen Inc., South Korea. The Basic Local Alignment and Searching Tool (BLAST) on the National Centre for Biotechnology Information (NCBI) website was used to confirm the presence of brugian parasites.

### Statistical Analysis

#### a. Calculating species diversity and abundance

The diversity of species in each mosquito collection was measured by the Shannon Diversity index (H) where the higher the value of H, the higher the diversity of species in the samples. The evenness or the abundance of species in each mosquito sample was measured by the Shannon Equitability Index (E_H_) (24).

#### b. Identification of dominant vectors

The vector Infection Rate (IR) and potential Infective Vector Density (IVD) were calculated as shown below to identify the dominant vector of BF in Sri Lanka.

#### c. Calculation of vector Infection Rate (IR) and potential Infective Vector Density (IVD)

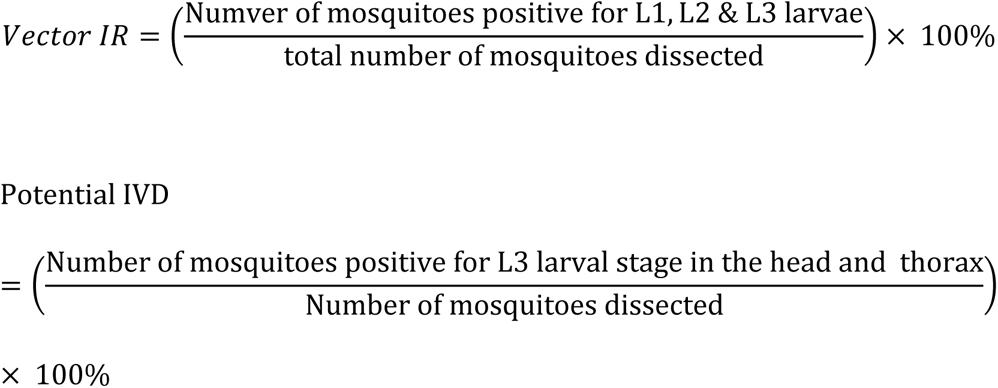

## Results

### Mosquito abundance and species diversity varies spatially and temporally

A total of 1711 mosquitoes from 19 species were captured **(Fig 3**). 72.5% (n= 1 245), 17.1% (n=293) and 10.4% (n= 178) mosquitoes were collected from dog-baited traps, window traps and gravid traps respectively. The highest number of species were collected from dog-baited traps, eighteen (18), while five (5) and ten (10) species were collected from gravid and window traps respectively.

**Fig 3:**
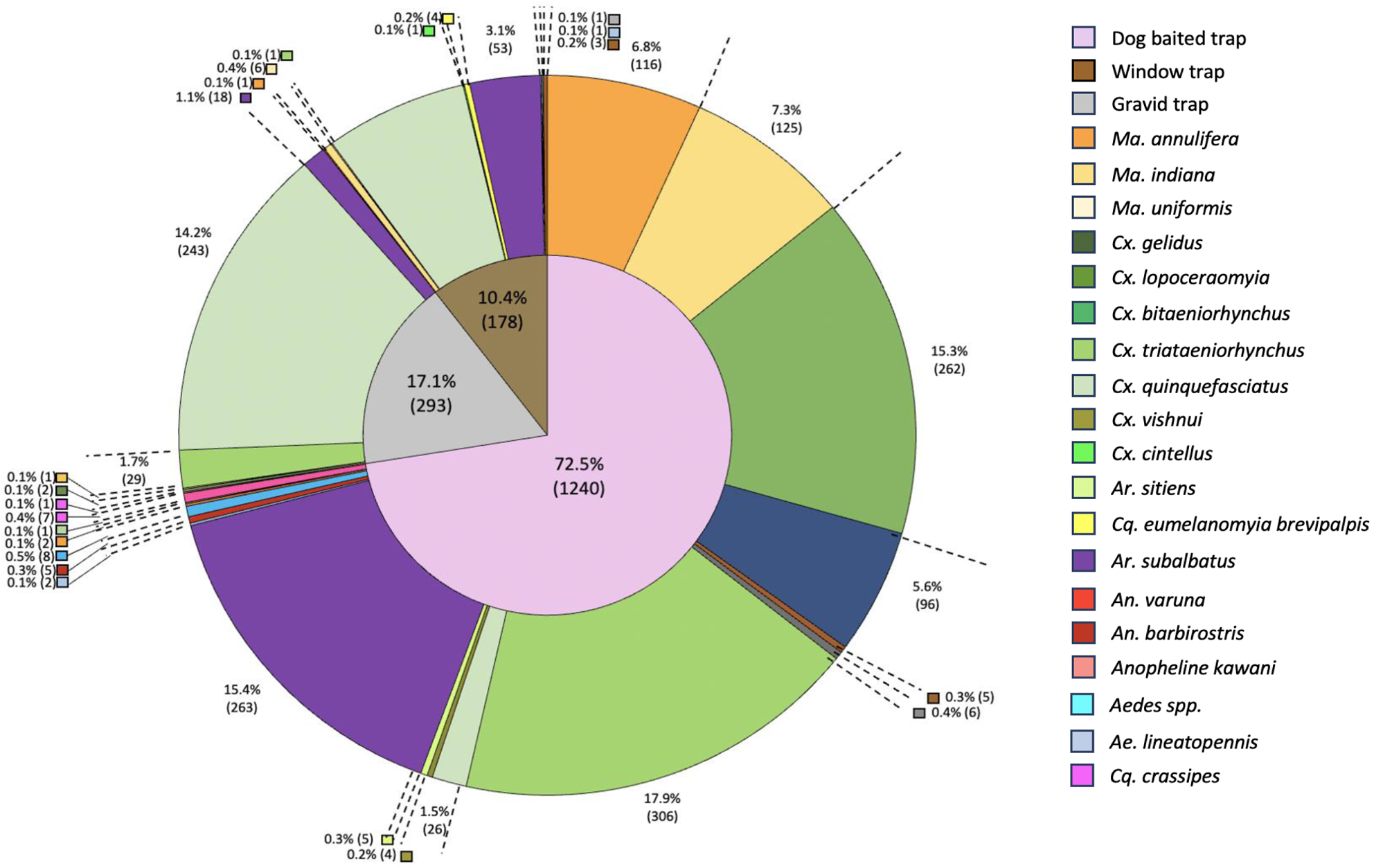
A species-wise representation of the total number of mosquitoes collected from each type of trap. A total of 1711 mosquitoes of 19 species were captured using dog baited, gravid and window traps.

The total mosquito abundance and species composition showed spatial and temporal variations between the study sites **(Fig 4)**. Mosquitoes were collected using varying numbers of trap settings (1 dog-baited trap, 1 window trap and 1 gravid trap) based on logistics at each study site and therefore, data was normalized to an average value per trap setting per site. S_1_ and S_2_ had the highest total mosquito abundance per trap setting (∼335) followed by L_3_ (195). The total mosquito abundance varied (S_1_: 254 vs 66, S_2_: 135 vs 208, S_3_: 168 vs 27) within each study site temporally; incidentally, the samples were one each from the monsoon and inter-monsoon seasons respectively. Regardless of the season, the highest mosquito species diversity was reported from the two sites in Puttalam, S_1_ (n=11, H_monsoon_ =2.322, H_intermonsoon_ =1.88) and S_6_ (n=08, H_monsoon_ =1.28). Four to six species were found in S_3_ (H_monsoon_ =1.27, H_intermonsoon_ =1.44), S_2_ (H_monsoon_ =0.64, H_intermonsoon_ =1.40) and S_4_ (H_monsoon_ =1.49), while only *Cx. quinquefasciatus* and *Ar. subalbatus* were found in the sample from S_5_ (H_monsoon_ =0.48). A high number of *Ma. indiana* was observed in S_1_, and to our knowledge this is the first record of a large number of *Ma. indiana* mosquitoes being reported in a catch in Sri Lanka.

**Fig 4:**
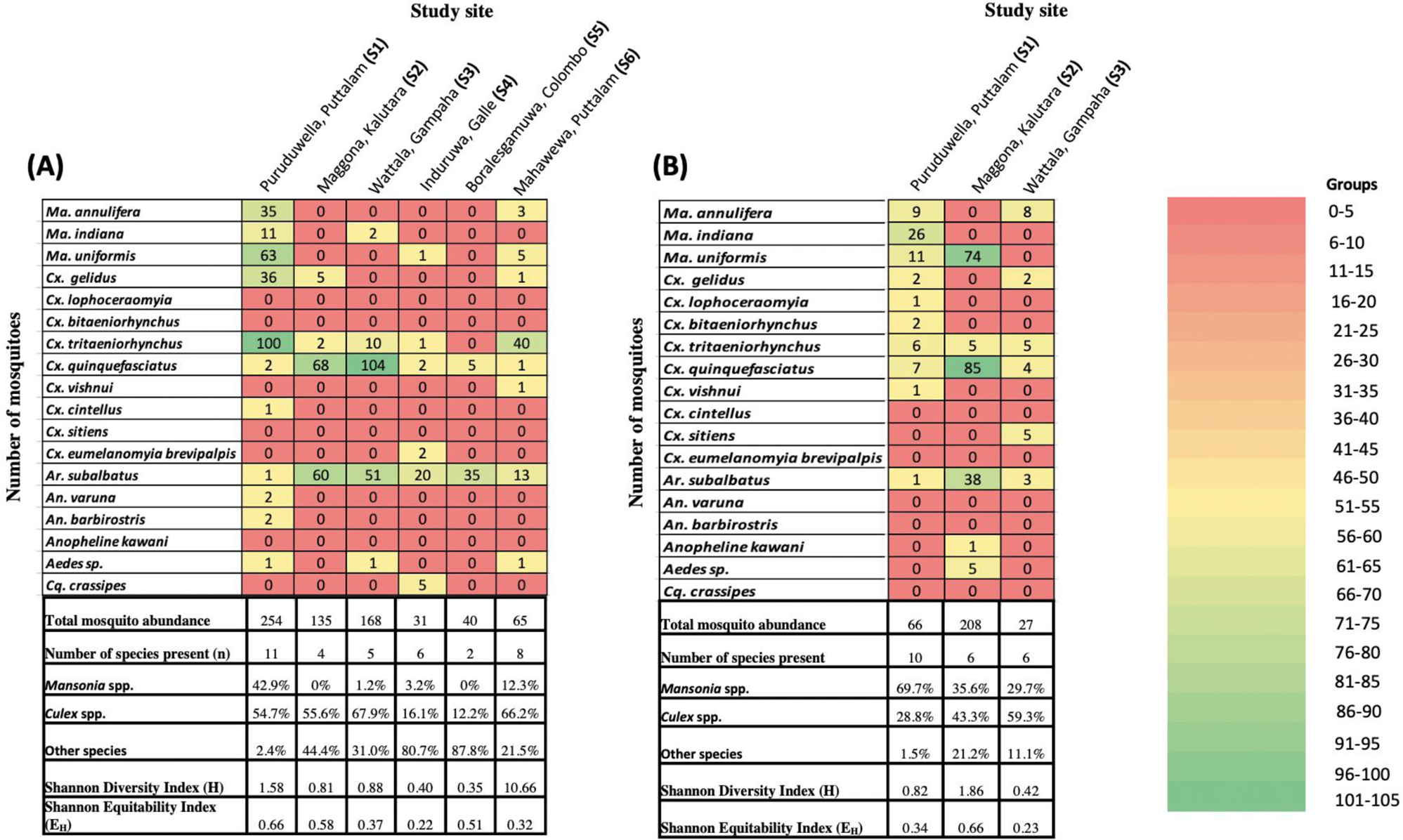
Mosquito abundance and species distribution varied spatially and temporally. Mosquitoes were collected using varying numbers of trap settings (1 dog-baited trap, 1 window trap and 1 gravid trap) based on logistics at each study site and therefore, an average value per trap setting per site within **(A)** monsoon and **(B)** inter-monsoon seasons is shown here.

The relative abundance of vector mosquitoes and the most abundant mosquito genus varied according to the geographical region and between seasons within the same region. *Culex* spp. was the most abundant in S_1_ in the monsoon and *Mansonia* spp. in the inter-monsoon. *Culex* spp. and *Armigeres* spp. were the most abundant in S_2_ in the sample from the monsoon season, while *Culex* spp. and *Mansonia* spp. were the most abundant in the inter-monsoon. *Culex* spp. mosquitoes were abundant in S_3_ during the monsoon, with the overall abundance of mosquitoes decreasing during the inter-monsoon and no species significantly abundant during that season. *Armigeres* spp. in the S_4_ and S_5_ and *Culex* spp. in S_6_ were the most abundant **(Fig 4).**

### Novel potential vector species identified

The presence of the L_3_ larval stage brugian parasites within the head and thorax regions of *Ma. annulifera, Ma. indiana, Ma. uniformis, Cx. tritaeniorhynchus, Cx. bitaeniorhynchus, Cx. quinquefasciatus*, *Cx. gelidus, Cx. lopoceraomyia, Cx. vishnui, Ar. subalbatus*, *Aedes spp.* and *Cq. crassipes* mosquitoes were subsequently molecularly confirmed. To our knowledge, this is the first report of *Ma. indiana, Cx. tritaeniorhynchus, Cx. bitaeniorhynchus, Cx. quinquefasciatus, Cx. gelidus, Cx. lopoceraomyia, Cx. vishnui, Ar. subalbatus, Aedes spp*. and *Cq. crassipes* with brugian larvae reported in the head and thorax regions in Sri Lanka, and *Cx. bitaeniorhynchus, Cx. quinquefasciatus, Cx. gelidus, Cx. lopoceraomyia* and *Cx. vishnui* in the world. This is suggestive of being potentially infective, thus a potential of serving as vectors of BF in Sri Lanka. Infection and potentially infective rates of mosquitoes varied between the study sites, and with the season within each study site. As shown in **fig 5**, the highest number of infected mosquitoes and the highest potentially infective vector density were observed from S_1_ and S_6_ study areas in Puttalam district, which incidentally is the district with the highest disease incidence.

**Fig 5:**
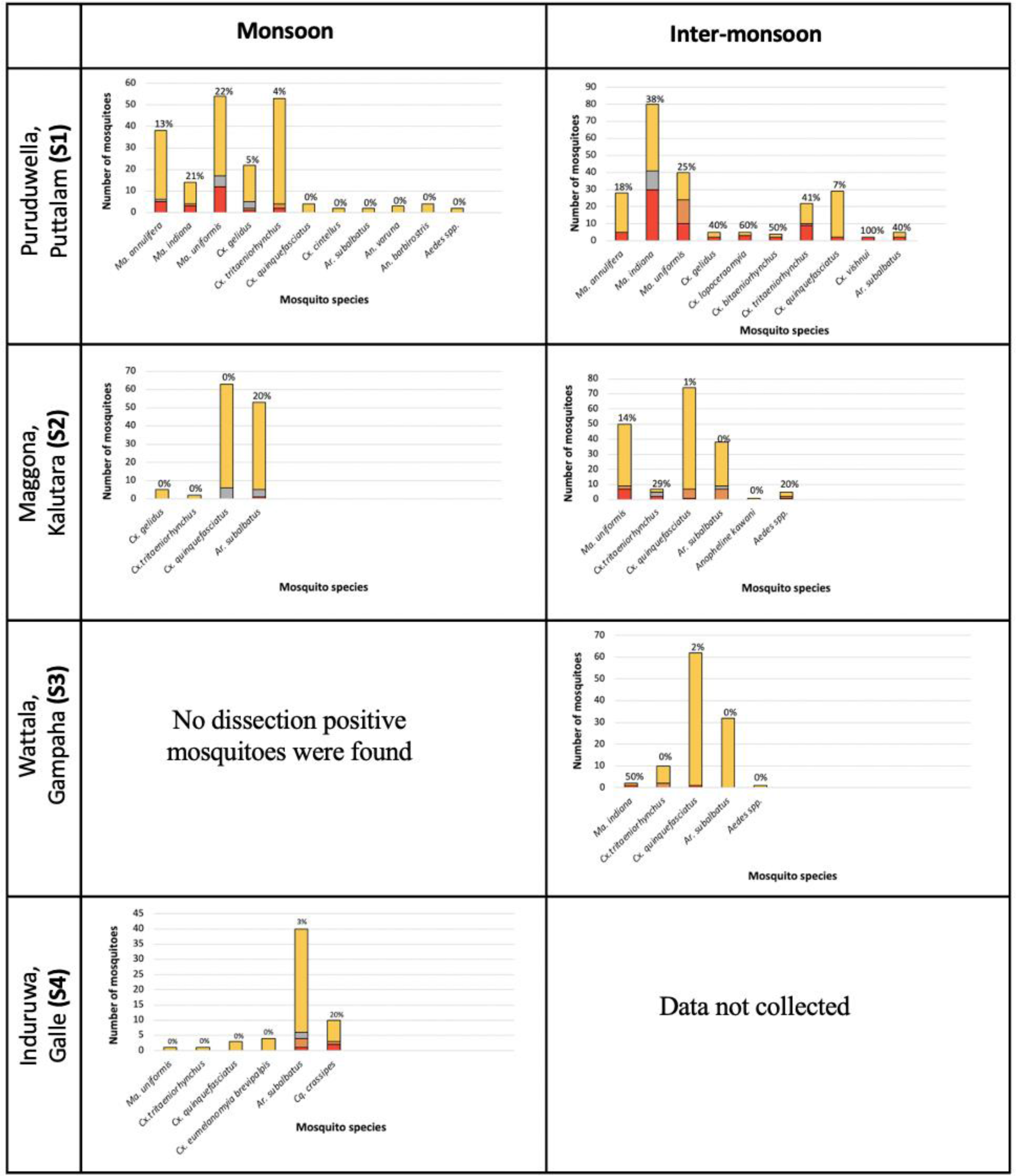
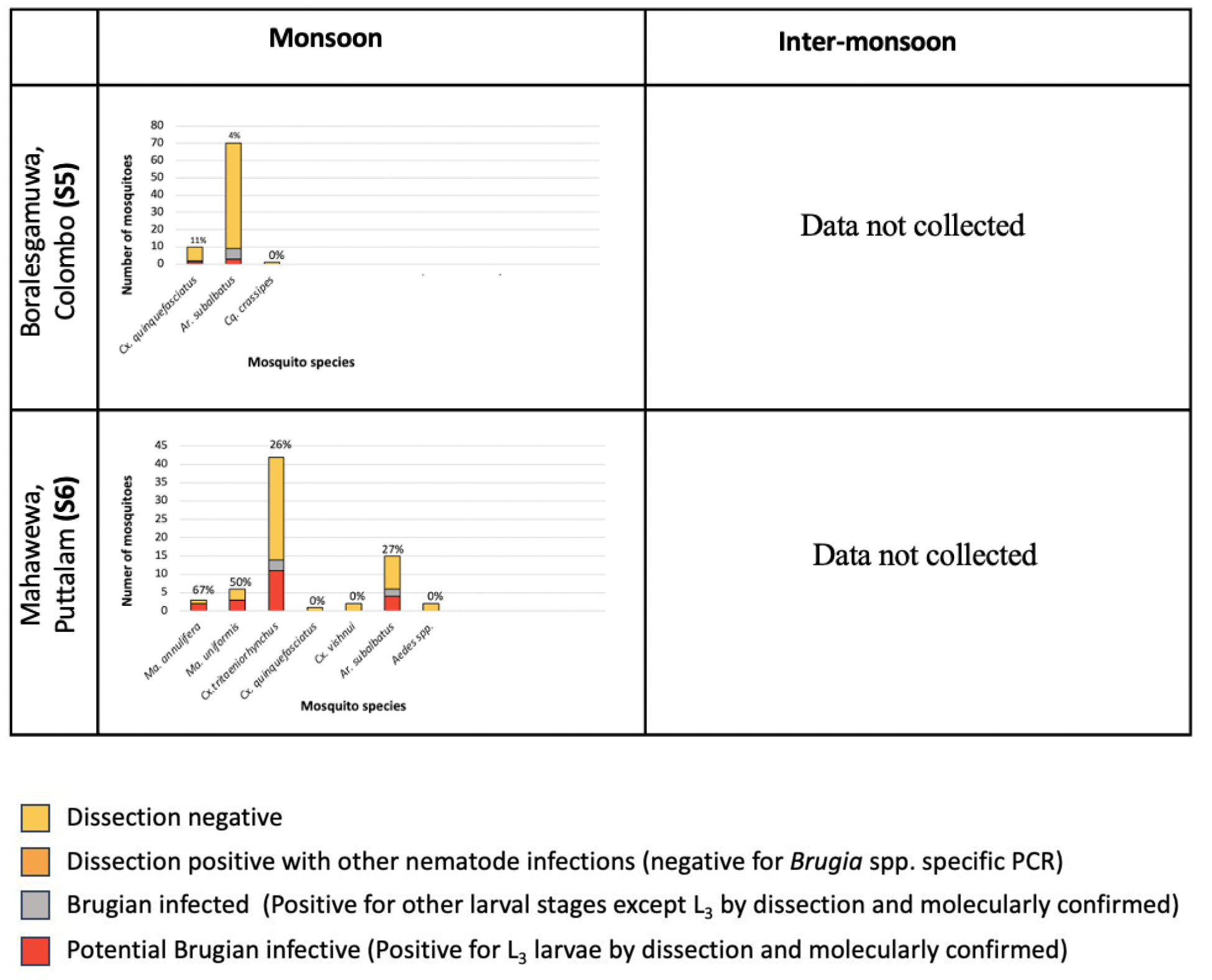
The abundance and species distribution of potential infective mosquitoes varied spatially and temporally. The head and thorax regions of a proportion of all the mosquito species from different geographical areas and seasons was dissected to identify mosquitoes with *B. malayi* infective L_3_ larval stage. Parasite positive mosquitoes were molecularly confirmed for BF-infections. Potential infective mosquitoes with parasites of L_3_ larval stage in the dissected head and thorax region of the mosquito and molecularly confirmed for brugian infections are represented as a percentage (percentage of potentially infective mosquitoes) for each species on the bars.

### Calculation of Infection Rate and potential Infective Vector Density

The mosquito species involved in active transmission of the infection varied between sites and with the season within each site as indicated by the Infection Rate (IR) and potentially Infective Vector Density (IVD) calculated for each mosquito sample (**Fig 6)**. The IR is a representation of the proportion of infected mosquitoes per species from the entire collection, while the potentially IVD provides some direction as to which mosquito species might potentially be involved in transmission of the infection at each site.

**Fig 6:**
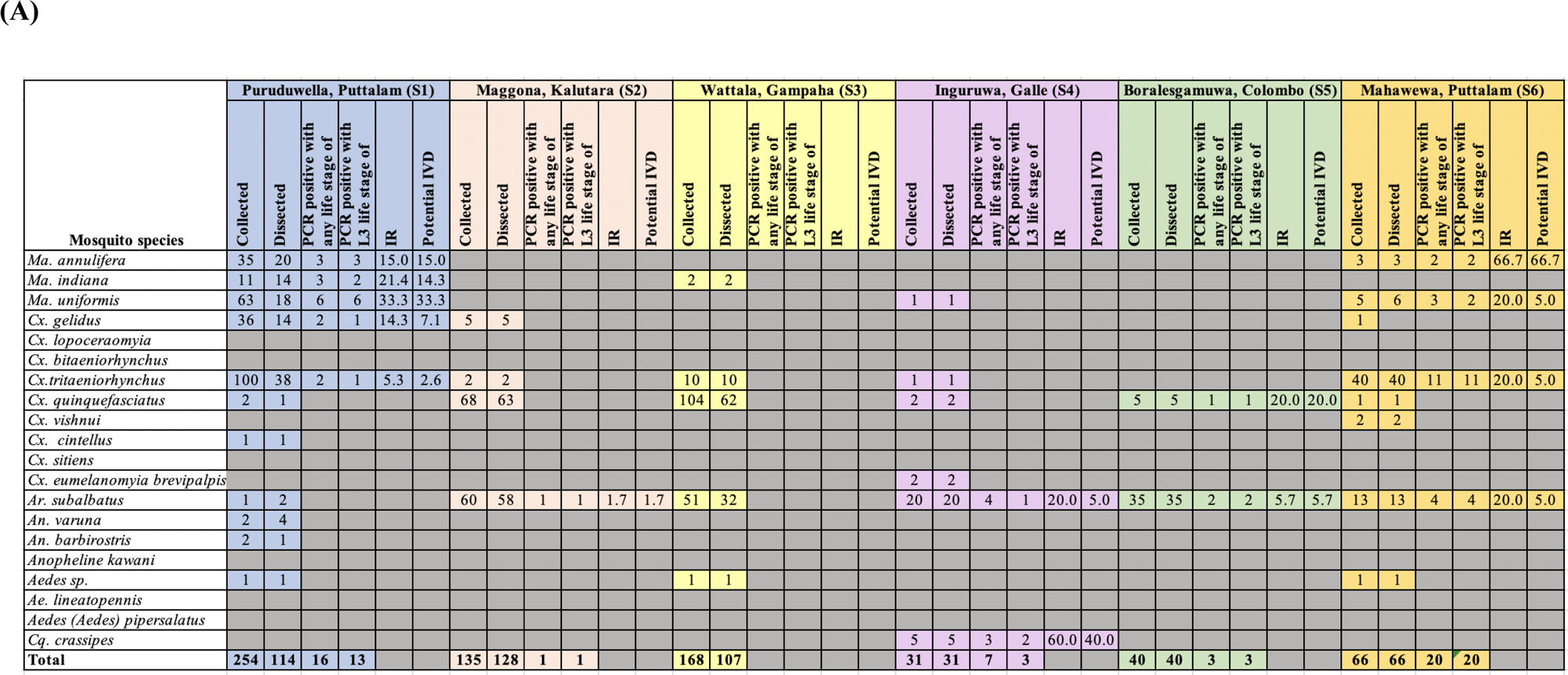
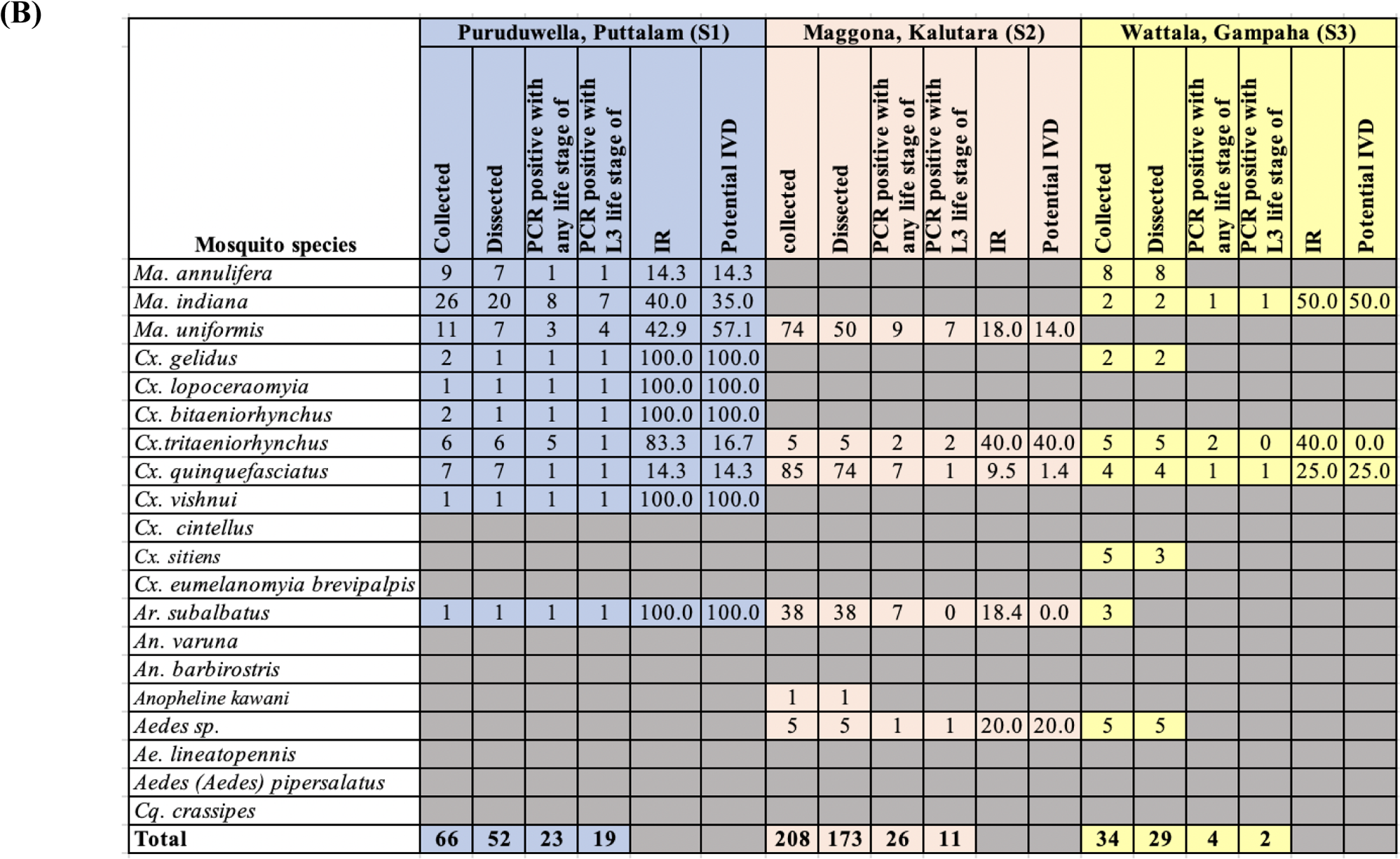
IR and VI calculated for all the mosquito species collected per trap setting, per day from the study areas. Cells in grey color represent mosquito species in which no parasites were observed. **(A)** monsoon and **(B)** inter-monsoon seasons are shown here.

During the monsoon season, the highest IRs were reported from *Mansonia* spp. mosquitoes from S_1_ and S_6_ sites. Number of infective mosquito species were high during the inter-monsoon period compared to monsoon in S_1_, S_2_ and S_3_ sites. Regardless of the season, S_1_ site had the highest risk of infection transmission.

### The abundance of infected vectors decreases with distance from the index site

The spatial distribution of infected vectors was investigated by comparing mosquito collections at varying distances within the study site at S_1_, where the most recent case was detected from the district with the highest disease incidence at the time of the study. A total of 281 female mosquitoes were collected at the index site. The number of female mosquitoes collected at 150 m site, 450 m site, and 650 m site was 224, 32, and 33 respectively **(Fig 7)** with abundance decreasing with distance from the index site. The most abundant mosquito species at the index site, 150m site, 450m site, and 650m sites were *Cx. tritaeniorynchus* (n=133, 47.3%), *Ma. uniformis* (n=89, 39.7%), *Cx. tritaeniorhynchus* (n=10, 31.3%), and *Ar. subalbatus* (n=24, 72.7%) respectively. 10 and 9 species were collected at the index and 150 m sites respectively and 6 species from the 450 m and 650 m sites. *Brugia* spp.-infected mosquitoes as confirmed molecularly were found only at the index site and the site 150 m from the index site with the number of infected mosquitoes decreasing with distance. Infected *Ma. indiana* and *Cx. tritaeniorhunchus* were present at the index site, infected *Cx. gelidus* at the 150 m site and *Ma. annulifera* and *Ma. uniformis* at both sites.

**Fig 7:**
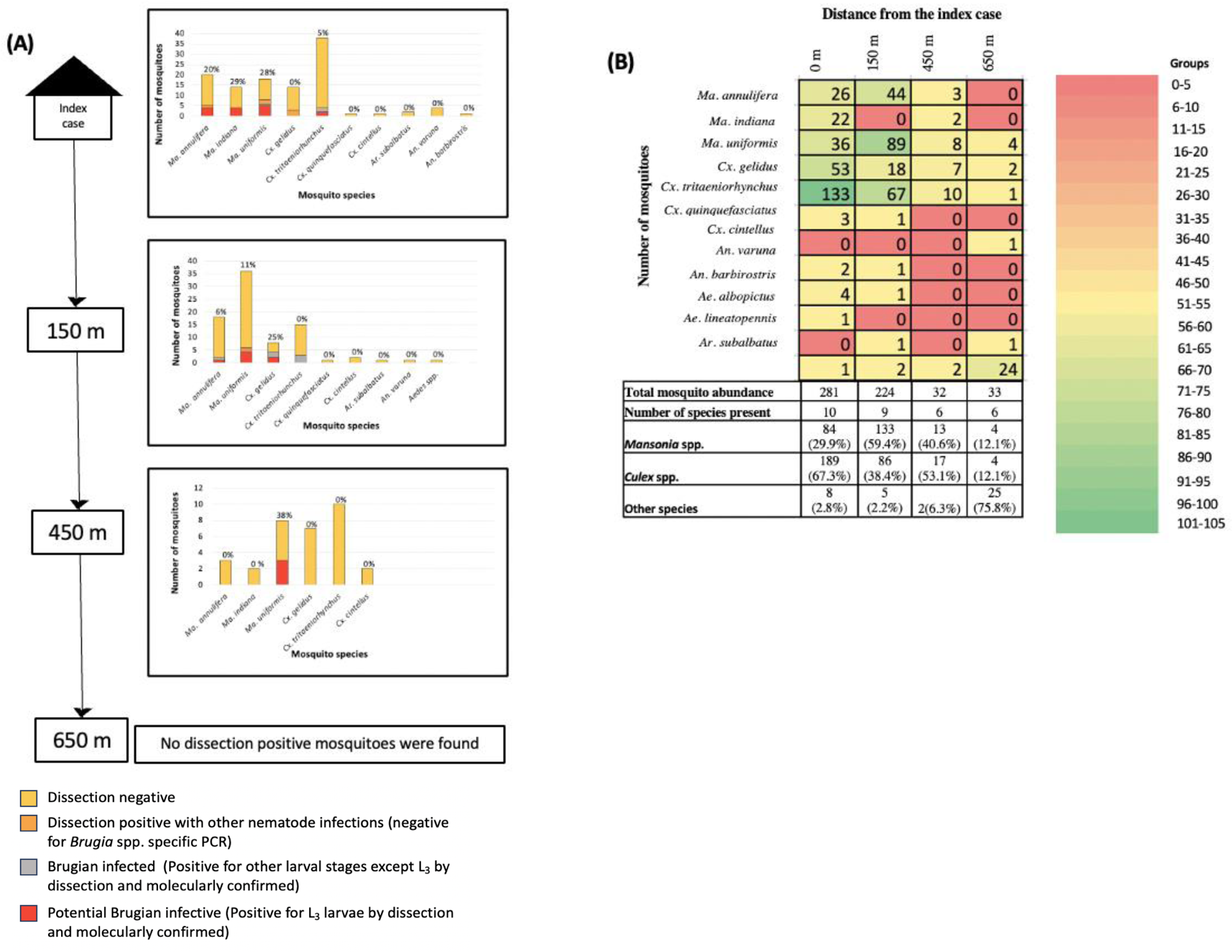
Species diversity and infectivity of mosquitoes decreased with distance from the study site. Spatial distribution of **(A)** total mosquitoes and **(B)** Brugian infective mosquitoes around a selected human BF index case were analyzed. The distribution was analyzed at the selected BF index case, 150 m, 450 m and 650 m away from the index case. A proportion of the mosquitoes from all the collected mosquito species were dissected for the detection of infectivity. The number of the potentially infective mosquitoes is represented as a percentage of the whole number of dissected mosquitoes on the bars.

## Discussion

Lymphatic filariasis (LF) is a debilitating disease that puts 859 million people at risk in 47 countries (25). Of the two forms of LF, Brugian filariasis (BF) can infect cats, dogs (20,26–30), monkeys (31) and pangolins (32) in addition to humans and therefore, can spread more rapidly than bancroftian filariasis. BF has reemerged in Sri Lanka after four decades of quiescence and show an increase in disease incidence in 2023. Lack of a cure for LF emphasizes the importance of prevention via effective vector control and Mass Drug Administration. This study was aimed at identifying the potential vectors of the newly emerged variant *B. malayi* parasite in Sri Lanka to aid in effective vector control via targeted measures.

Here we report evidence of infection and development of the *B. malayi* parasite in *Ma. annulifera, Ma. indiana, Ma. uniformis, Cx. tritaeniorhynchus, Cx. bitaeniorhynchus, Cx. quinquefasciatus*, *Cx. gelidus, Cx. lopoceraomyia, Cx. vishnui, Ar. subalbatus*, *Aedes spp.* and *Cq. crassipes* in Sri Lanka. To date, only *Ma. annulifera* and *Ma. uniformis* have been reported as potential vectors for BF in Sri Lanka (33–35). Although, detection of L_3_ larval stage brugian parasites within the head and thorax region of the mosquitoes, suggest that these mosquitoes might be able to transmit the disease, some studies have demonstrated the inability of some mosquitoes in transmitting the disease even so (36). Therefore, further studies need to be conducted to investigate the vectorial capacity of the potential vectors identified by the current study. Moreover, bancroftian filariasis cases have been reported to the AFC from these sites and dirofilarial cases have been detected in the canine population within the same area. Hence the dissection positive and PCR negative mosquitoes may be involved in the transmission of either bancroftian filariasis or dirofilariasis in this area.

Interestingly, *Ma. indiana* was by far the most prevalent and potentially infective mosquito species at the study site of the most recent human BF case at the time of the study, closely followed by *Ma. uniformis* and various *Culex* species; Incidentally this is also the district with the highest number of reported human BF cases to date. This is also the first study reporting a high number of *Ma. indiana* in Sri Lanka. Three (3) *Ma. indiana* mosquitoes were reported in a study conducted in the Gampaha district, which were not infected with brugian parasites (20) and none in a survey done for six consecutive months where nearly 7000 mosquitoes were analyzed (37). However, *Ma. indiana* has been reported in abundance in filariasis endemic areas in other countries (18) and has been reported as a potential vector for filariasis in other countries like Thailand, Malaysia and Indonesia (38). Studies have shown the capability of *Ma. indiana* in transmitting nocturnally sub-periodic *B. malayi* parasites (38), supporting its vector potential for BF. *Ar. subalbatus* has been reported as a vector for zoonotic *B. pahangi* in Thailand (39) and Malaysia (40,41), *Ae. albopictus* and *Cx. tritaeniorhynchus* as vectors of *B. malayi* in Indonesia (42) and *Cq. Crassipes* as a vector for brugian parasites in Malaysia (43).

This is the first report of *Ma. indiana*, *Cx. bitaeniorhynchus, Cx. quinquefasciatus*, *Cx. gelidus, Cx. lopoceraomyia* and *Cx. vishnui* as potential vectors of BF in Sri Lanka. In 1995 Bangs et al. have reported the susceptibility of *Culex tarsalis* and *Culex erythrothorax* to sub-periodic *B. malayi* by laboratory experiments and thereby have provided evidence for the ability of the genus *Culex* in acting as vectors for BF (44). Interestingly, certain laboratory experiments have shown the complete refractoriness of *Cx. sitiens* to sub-periodic *B. malayi* (45) and the mid gut acting as a barrier for the brugian filarial parasites in *Cx. pipiens pipiens* (46). However, a study done by Erickson *et al.* have reported the presence of filarial DNA within *Cx. pipiens* head region, providing evidence for the potential of *Cx. pipiens* acting as vectors for BF (47).

Abundance and composition of mosquito species varied between study sites in the current study. *Mansonia spp.* mosquitoes were found in higher numbers from both the study sites at Puttalam (Puruduwella and Mahawewa). The two study sites boarded a lake with aquatic plants, particularly floaters like water hyacinth, *Salviania* and *Pistia stratiotes*, which are well-known breeding sites for *Mansonia* spp. (48) (18). There were also paddy fields along one side of the Puruduwella study site, which could be the reason for the high number of *Cx. tritaeniorhynchus* at the site (49). *Cx. quinquefasciatus* were found in relatively higher densities in the Maggona-Kalutara where there was a water tank with mosquito larvae and another with water Hycinth, and at Wattala-Gampaha which is within an urban area and had a higher population density compared to other study sites. According to previous studies, *Cx. quinquefasciatus* mosquitoes thrive in obstructed drains and stagnant water bodies contaminated by human activities in sites with a higher human population. A higher abundance of *Armigeres* spp. was observed at Induruwa-Galle and Boralesganuwa-Colombo study sites which are in more urban settings when compared to the rural (Puruduwella) and semi-urban (Mahawewa) areas in Puttalam, an observation supported by literature (50).

Abundance and composition of mosquito species also varied between seasons at the same study site. In general, the mosquito population in Sri Lanka is more prolific during the monsoon season than the inter-monsoon season (49,51). The rainy season creates ideal conditions for maximizing the vector Infection Rate (IR), which is based on factors such as rapid parasitic development, proper nursing, and low parasitic damage or mortality (52). As expected, a higher mosquito density was observed during the monsoon period except in Maggona-Kalutara. The probable reason for the anomaly in Kalutara is the recent clearance of aquatic vegetation near the study site at the time of sampling. Interestingly, the number of infective mosquitoes was higher during the inter-monsoon period in all the three sites studied during both seasons. Chandra *et al.* reports a lack of coordination between the transmission season and the period of highest vector density where high filarial transmission was found to occur during the hot months of the rainy season and sometimes in summer (53). This lack of synchronization hinders the transmission process and keeps it at a relatively low level (53). They state that this lack of coordination may pave the path to naturally limit the BF transmission within the country and is applicable to the transmission pattern in Sri Lanka as well. The seasonal variation was examined only in the three study sites from the districts with the highest number of BF cases reported and therefore, needs to be extended to other regions of the country for a comprehensive understanding.

Many factors such as Vector Index (VI), Infectivity Rate, vector potency, flight capacity of infected mosquitoes and number of vector-host encounters determine the rate of transmission of vector-borne diseases (54). Analyses of the spatial distribution of mosquitoes at the study site with the highest abundance revealed that infective mosquitoes were detected within a 150 m, 450 m and 650 m radius from the index site although the potential vector species were still abundant up to about 600 m from the index site. And the number of potential vector mosquito species reduced with the distance from the index case. This can be due to the reduced flight capacity of mosquitoes upon parasitic infection (55). Mosquitoes with developing L_1_ and L_2_ larvae have shown a reduced activation and flight ability, while L_3_ infective mosquitoes have shown an increased activation and flight towards the host (56). Furthermore, infection of *Ae. aegypti* Liverpool (LVP) strain mosquitoes with *B. malayi* have shown a decreased flight speed compared to uninfected mosquitoes. The detrimental effect of parasitic infection on flight capacity is due to the additional energy cost that the mosquito must spend to support the development of the parasite (57). Therefore, the differences in the flight capacity of potential vectors upon parasitic infection can affect the spatial distribution of vectors and thereby the ability of vectors to transmit the disease. Collectively, *Ma. indiana, Ma. uniformis, Ar. subalbatus*, *Cx. tritaeniorhynchus* and *Cx. quinquefasciatus* have the highest vector indices. The recovery of parasite-positive *Ma. indiana, Cx. quinquefasciatus* and *Ar. subalbatus* mosquitoes from window traps demonstrates the probable anthropophilic nature of these species and their high prevalence in the respective areas deems further investigation on their vector potential essential.

This is the first study reporting the presence of brugian parasites in multiple genera of mosquitoes in Sri Lanka. This is also the first report in the world of the presence of these parasites in *Cx. bitaeniorhynchus*, *Cx. gelidus, Cx. lopoceraomyia and Cx. vishnu.* Ughasi *et al*. reported the possibility of mosquito species previously considered as non-vectors to act as vectors of filariasis parasites over generations (58). This may reflect a parasite with genetic modifications and higher pathogenicity which needs further investigations. Data generated by the current study could be employed to refine prevention and control strategies by developing more tailored vector control measures in Sri Lanka and other BF endemic countries.

**Table 1:**
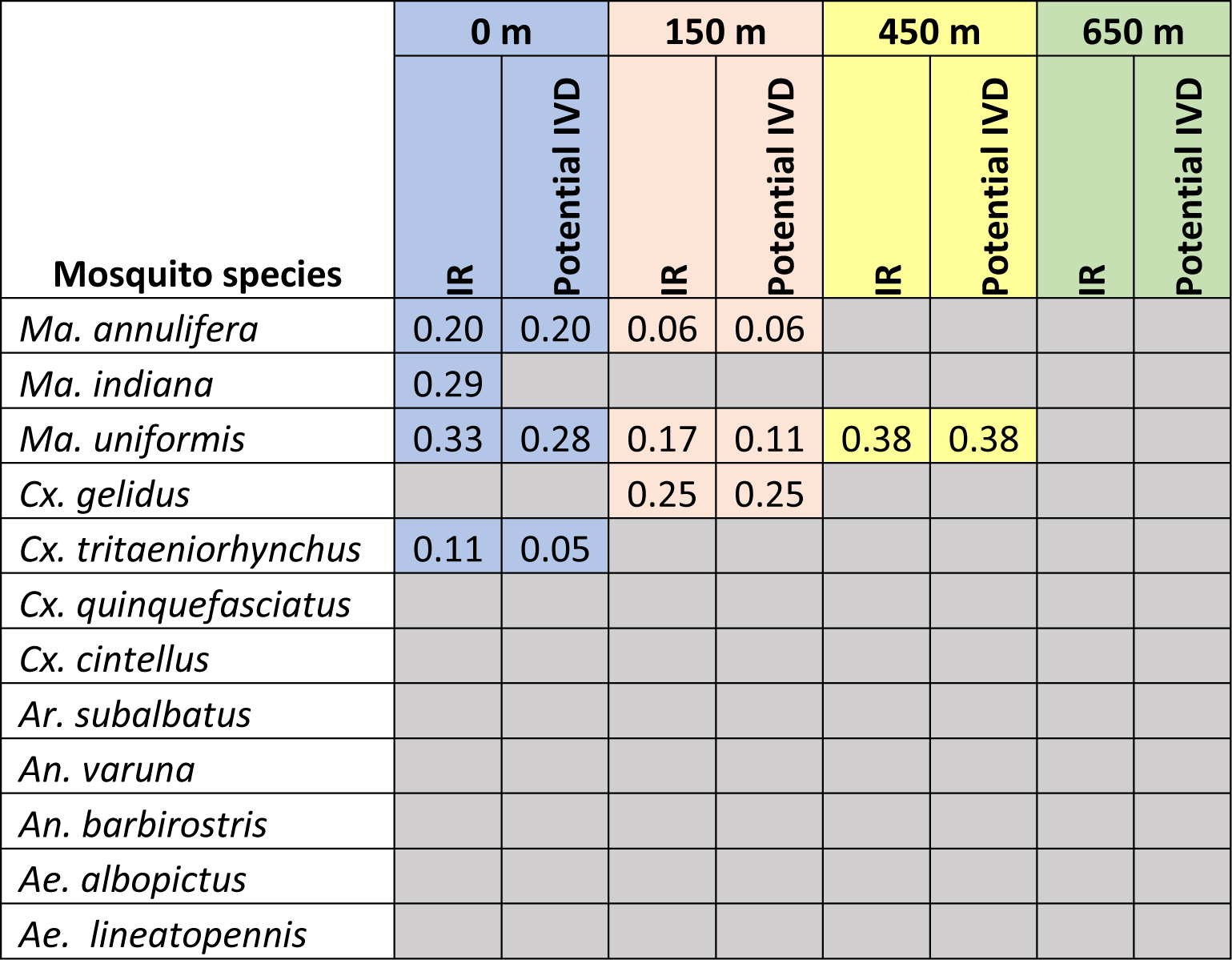
The Infection Rate (IR) and potentially Infective Vector Density (IVD) deferred with distance within the same site. . IR and potentially IVD calculated for all the mosquito species collected from the index case, 150 m, 450 m and 650 m away from the index case. Cells in grey color represent the mosquito species in which no parasites were observed.

## Acknowledgments

The authors wish to acknowledge the support extended by the staff of the Anti-filariasis Campaign, Colombo, Sri Lanka, the staff of Regional Director of Health Services branches at Madampe, Colombo, Kalutara and Galle, Sri Lanka and the staff of the Department of Entomology at the Medical Research Institute, Colombo, Sri Lanka and Mr. Lahiru Herath. The authors also wish to acknowledge the support rendered by Dr. Dilakshini Dayananda of the Genetics and Molecular Biology Unit, University of Sri Jayewardenepura, Sri Lanka and Dr. Sachini Fernando of the Center for Biotechnology, University of Sri Jayewardenepura, Sri Lanka for data analysis and reviewing the manuscript.

## Notes

### Competing Interest Statement

The authors have declared no competing interest.

